# Interactions between weather, ungulates and geese create opportunities for wood-pasture cycles

**DOI:** 10.1101/2024.07.19.604241

**Authors:** Koen Kramer, Perry Cornelissen, Geert W.T.A. Groot Bruinderink, Loek Kuiters, Dennis Lammertsma, J. Theo Vulink, Sip E. van Wieren, Herbert H.T. Prins

## Abstract

Managing nature reserves to achieve coexistence of ungulate species and vegetation heterogeneity is challenging in small reserves. A central question is whether spatial heterogeneity is maintained or restored through occasional natural disturbances. This is the premise of the wood-pasture hypothesis, which asserts that natural disturbances, such as winter severity, are sufficiently significant to sustain the long-term coexistence of ungulate species and provide opportunities for the establishment of shrubs and trees, thereby enhancing vegetation heterogeneity. Testing the wood-pasture hypothesis under field conditions necessitates long-term time series of both vegetation and herbivore dynamics. Therefore, we employed a modeling approach to test this hypothesis.

The study area was the Oostvaardersplassen, a nature reserve in the Netherlands characterized by rich and productive clay soil. This area hosts an assemblage of Heck cattle, Konik horses, and red deer. Additionally, large numbers of geese frequent this reserve, competing for the same food resources. The Oostvaardersplassen is one of the few areas in Europe where neither the populations of coexisting herbivore species nor the vegetation have been managed and its development was monitored for several decades. This provides unique insights into natural herbivore-vegetation interactions that can be applied to the management of other nature reserves.

The results indicate that weather variability and the presence of both geese and large ungulates are essential factors in creating a heterogeneous landscape. The general pattern is that weather variability induces fluctuations in ungulate numbers, and geese reduce the numbers of ungulates. Coexistence of the three ungulate species remains possible regardless of weather variability and the presence of geese. However, the presence of the largest ungulate, cattle, is threatened when weather is highly variable and geese numbers are high. The vegetation is likely evolving towards a predominantly open landscape with occasional opportunities for the establishment of spiny shrub species. Nature reserves managed with an assemblage of ungulate species, which numbers are controlled by food supply, winter severity and competition, should thus focus on the substantial presence of a few species (“biomassality,” e.g., for migratory birds) rather than achieving the highest species diversity in a small area (“biodiversity”).

## Introduction

Inspired by contemporary natural or near-natural grazing systems in Africa and North America, as well as by past Pleistocene and Holocene ecosystems, the (re-)introduction of wild ungulates has garnered significant attention (Caro & Sherman, 2009; Hodder & Bullock, 2009; Huynh, 2011; Jackson & Hobbs, 2009; Navarro & Pereira, 2012; Rey Benayas et al., 2009; Svenning, 2002). This management strategy aims to restore ecosystem functionality. In Western Europe, alongside wild ungulates such as European bison and red deer, domestic cattle and horses have been introduced as substitutes for their wild ancestors (Hodder & Bullock, 2009; Kirby et al., 1995; Olff et al., 1999b; Vera, 2000; WallisDeVries et al., 1998). In fragmented landscapes, these wild and domestic ungulates are introduced in small fenced areas without large predators (Lindenmayer & Fisher, 2006). Under these conditions, ungulate populations are primarily regulated by food supply and winter conditions (Coulson et al., 2001).

Although this management system has been practiced for the last 30 years in some European countries, little is known about the long-term population dynamics of the ungulates and their effects on vegetation (McCann, 2007). We analysed long-term population dynamics of ungulates and their effects on vegetation using a process-based model. This model was applied to one of the first areas in Europe where a self-regulating, multi-species assemblage of ungulates was introduced: the Oostvaardersplassen nature reserve in the Netherlands.

In the eutrophic wetland of the Oostvaardersplassen (OVP, 5,600 ha), an assemblage of cattle, horses, and red deer was introduced in the 1980s. The area is fenced, and populations were not controlled at fixed stocking rates until 2018. Until that year, individuals that had no chance of survival were culled to prevent unnecessary suffering, no supplementary feeding was provided, and large predators were absent. Additionally, the reserve was visited annually by thousands to tens of thousands of Greylag and Barnacle geese. A few years after their introduction, the populations of the ungulates grew exponentially. After these initial years, the growth rate stabilized, and the numbers reached a maximum. After 2018, the management has changed and animal numbers are controlled to much lower densities.

Corresponding with the increased herbivore population, the vegetation transitioned from a heterogeneous mixture of grasslands, tall herbs, reed, scrub, and trees to a homogeneous vegetation dominated by short grazed grasslands (Cornelissen et al., 2014a). Over the last 10 years, the cattle population has decreased, while the populations of horses and red deer, as well as the total number of geese, have continued to increase. As the vegetation becomes dominated by short-grazed grass species, competition among the different ungulate species intensifies (Cornelissen & Vulink, 2015; Menard et al., 2002; Putman, 1996). This competition could lead to the exclusion of the less competitive species. In this scenario, cattle could be outcompeted by other ungulates and geese, as cattle cannot graze on short swards as efficiently as the other herbivores can (Menard et al., 2002).

Even in scenarios of intense competition for food, mechanisms exist that enable ungulates to coexist. For instance, in heterogeneous areas with alternative forage sources, one species can alter its diet and habitat use to exploit these alternatives (Putman, 1996). In the Oostvaardersplassen (OVP), this mechanism is evident as horses dig into the soil to consume roots of Phragmites and Urtica, which are inaccessible to other herbivore species. Another mechanism promoting coexistence involves occasional disturbances such as climatic variations, predation, pests, and diseases (Coulson et al., 2001; Hopcraft et al., 2010; Sinclair et al., 2003). These disturbances temporarily reduce the populations of all ungulates or the most dominant competitor, thus preventing competitive exclusion. A reduction in ungulate numbers leads to an increase in per capita forage availability, allowing competing species to coexist.

A reduction in ungulate numbers is a crucial aspect of the wood-pasture hypothesis (Olff et al., 1999b; Vera, 2000). This hypothesis posits that high ungulate densities facilitate the transition from woodland to grassland through browsing and bark stripping, which causes shrub and tree mortality (Gill, 2006). Concurrently, ungulates maintain short-grazed grasslands, creating opportunities for shrubs and trees to re-establish in these natural pastures. Such re-establishment requires a temporary reduction in herbivore densities for a sufficient duration (Cornelissen et al., 2014; Olff et al., 1999a; Schippers et al., 2014; Vera, 2000). Variability in weather conditions or in the numbers of small herbivores, such as geese, can induce fluctuations in ungulate populations, providing windows of opportunity for woody plant species to establish. If the reduction in herbivore populations is significant, woody plant species can grow to a height where their foliage becomes inaccessible to herbivores, thus escaping being browsed.

In this study, we assessed the effects of variability in weather conditions and in the numbers of a small herbivore, namely geese, on the dynamics of ungulates and vegetation heterogeneity. We utilized the model FORSPACE (Kramer et al., 2006; Kramer et al., 2003), long-term observational weather data, and historical records of visiting geese. The research questions addressed in this study are:

A. Is weather variability sufficient to prevent competitive exclusion of the ungulates, allowing them to coexist?
B. Can geese reduce ungulate numbers through competition by grazing the sward to even lower heights than the ungulates?
C. Does a temporary reduction in ungulate numbers provide windows of opportunity for woody species to establish?

## Material and Methods

### Research area

The OVP area (52°26’ N, 5°19’ E) is a eutrophic wetland spanning approximately 5,600 hectares in the Zuidelijk Flevoland polder of the Netherlands, reclaimed from Lake IJsselmeer in 1968. As a former lake, the soil comprises 30-35% clay. Three distinct habitat types characterize the research area: grasslands (dominated by *Agrostis stolonifera* L., *Lolium perenne* L., and *Trifolium repens* L.), reed vegetation (*Phragmites australis* (Cav.) Steud.), and a semi-open mosaic of reed, tall herbs (*Urtica dioica* L., *Cirsium* spp. Mill.), elder (*Sambucus nigra* L.), and willow (*Salix* spp.) (Jans and Drost, 1995). Most willow species, primarily white willow (*Salix alba* L.), established on bare soil in 1968-1969 after the area was drained and dried. Elder established several years later, from the early 1970s until the early 1990s. Elder produces cyanogenic glucosides (Atkinson and Atkinson, 2002), which can be toxic or lethal (Majak and Hale, 2001). Ruminants are better equipped to counteract these toxic compounds than hindgut fermenters (Van Soest, 1994), as demonstrated by Vulink (2001) for the Oostvaardersplassen.

Cattle, horses, and red deer were introduced to the OVP area in different years: 32 Heck cattle (*Bos taurus* L.) in 1983, 18 Konik horses (*Equus caballus* L.) in 1984, and 52 red deer (*Cervus elaphus* L.) in 1992. By January 2015, the populations had grown to approximately 250 cattle, 1,200 horses, and 3,200 red deer. The ungulate populations were counted annually. Prevalent geese species in the OVP area are Greylag geese (*Anser anser* L.) and Barnacle geese (*Branta leucopsis* L.), which were counted weekly along a fixed route through the grasslands from 1996 to 2014. The annual average number of geese increased from about 3,000 per day in 1996 to about 10,000 per day in 2014. Both species are present year-round, with peak numbers during winter and spring.

### Model description

We utilized the spatially explicit process-based model FORSPACE to delineate the interactions between vegetation development and herbivore density (Kramer et al., 2006; Kramer et al., 2003) (Fig. 1). In this model, plant populations are defined by plant density, the biomass of various plant components, and their structural attributes. These variables are computed for each tree, shrub, herb, or grass species. Ungulate populations are described by the weight and number of both juvenile and adult cohorts for each species. The model operates on a monthly time step with a spatial resolution of 1 hectare for the OVP area. Comprehensive model descriptions, including sensitivity analyses and validations for other areas, can be found in Kramer et al. (2006), Kramer et al. (2003), and Kramer (2001). The model is implemented in the dynamic GIS software PC-Raster (Wesseling, 1996).

**Fig. 1.**
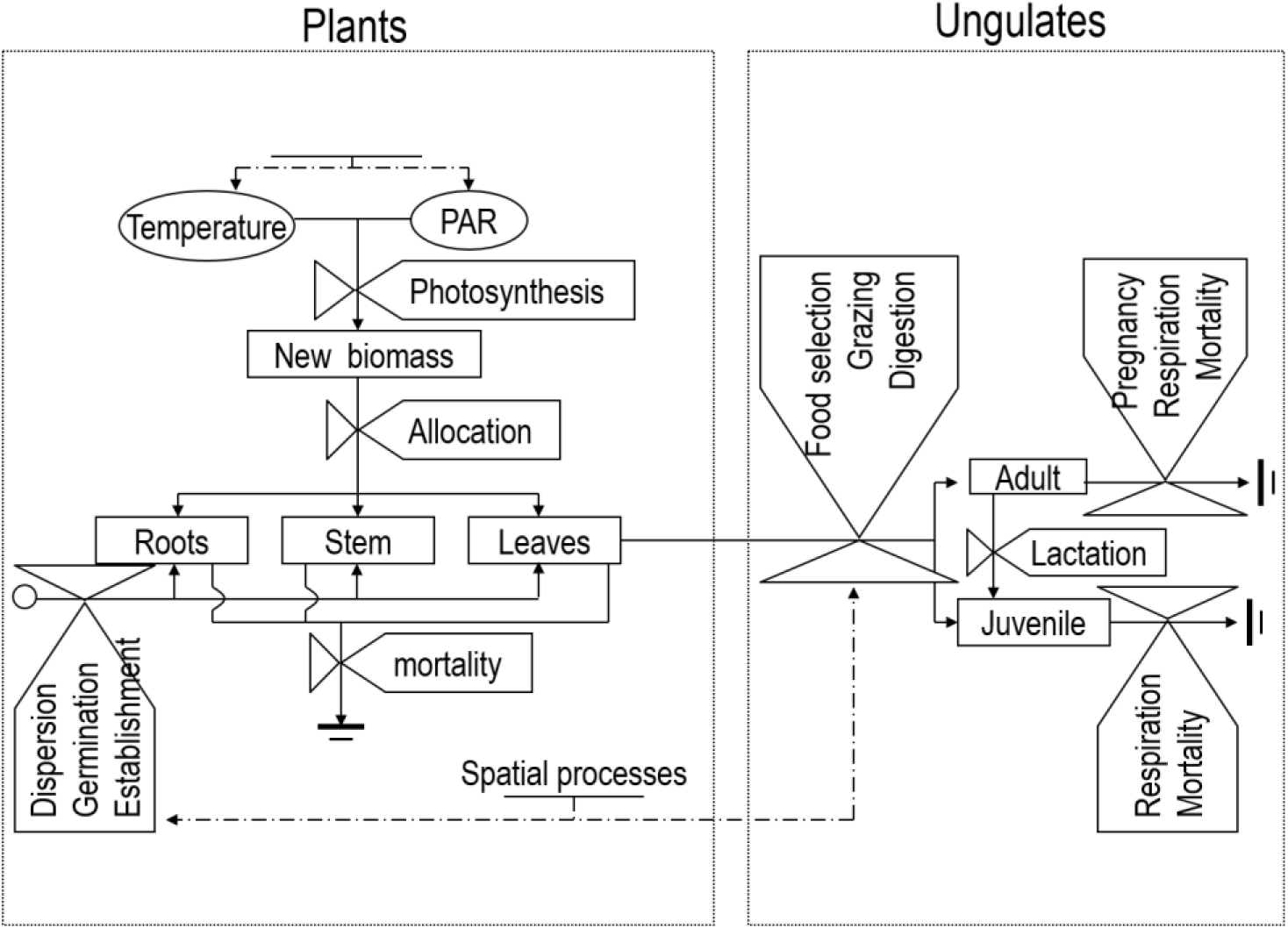
Flow diagram showing the principal processes and flow of information between plants and ungulates in the model FORSPACE.

Below, we outline the necessary model adjustments to address the specific questions of this study: the impact of snow cover on food availability, the inclusion of geese as small herbivores, grazing by geese, parametrization of willow and elder for fertile soils, the addition of hawthorn as a new species, and the effect of low winter temperatures on the survival of ungulates.

The model adjustments address the following issues:

i. *Weather Data*: We utilized weather data from station De Bilt of the Royal Dutch Meteorological Institute (KNMI), covering the past 110 years, as data collection began in 1901. This station is situated 38 km from the OVP area. Snow cover is defined as the fraction of the month during which vegetation is covered by snow. It is assumed that herbivores cannot access plants when snow cover is present.
ii. *Geese Populations*: Large numbers of moulting and wintering geese visit the OVP area year-round, consuming a substantial portion of the annual net primary production. The vegetation intake by geese is modeled similarly to that of ungulates. However, the population dynamics of geese are not simulated, as their population is primarily influenced by external factors such as food availability outside the OVP area (Van Eerden, 1998). During days with snow cover or mean daily temperatures below 0°C, the number of visiting geese is set to zero, as they migrate to warmer areas without snow cover.
iii. *Woody Species Parameters*: The parameter values for willow and elder were adjusted based on measurements of growth and development of these species in the OVP area (Cornelissen et al., 2014a, b).
iv. *Thorny Shrubs*: The plant functional type ‘thorny shrubs’ was added, exemplified by hawthorn (*Crataegus monogyna* Jacq). All ungulates induce hawthorn mortality by bark peeling. This effect is incorporated into the model by increasing the turnover of hawthorn individuals if the total number of herbivores exceeds 800 animals, comparable to the total number of ungulates in 1996. Before 1996, new establishments of woody species were visible in aerial photographs (Cornelissen et al., 2014b), but after this year, no new establishments were seen, and no seedlings of woody species were found in the field (Cornelissen et al., 2014a, b). We define the presence of thorny shrubs as hawthorn and palatable shrubs as willow and elder exceeding 1.5 m in height. The model assumes that from this height onward, ungulates do not affect the height of thorny shrubs. However, the height of other shrubs can be reduced until the plant exceeds the herbivore-specific maximum browsing height. This differentiation is made because hawthorn can develop a central shoot that escapes browsing, a process not present in willow or elder.
v. *Winter Temperature Effects*: The effect of winter temperature on ungulate survival is simulated by increasing their maintenance costs if the average monthly temperature drops below 0°C. Due to limited information in the literature on the enhancement of maintenance respiration with decreasing freezing temperatures, the model was calibrated under the assumption that each herbivore species in the model survived the most severe winter between 1901 and 2013. This assumption is supported by Harris et al. (1995), who observed significant mortality among ungulate species during severe winters but no local extinctions. Additionally, horse populations in Mongolia experience severe winter losses, yet some animals survive (Kaczensky et al., 2011). A similar survival pattern is observed in reindeer populations (Klein, 1968). An extensive literature review found no evidence of wild ungulate populations going extinct due to adverse winter conditions in any large nature reserve in Northwest Europe during the period 1901– 2013.

### Model validation

A model versus data comparison was performed between the observed and predicted number of herbivores over the period 1996-2013, based on the modifications described above. We used 1996 as a starting point because from that year, the entire area was grazed by cattle, horses, and red deer. The actual number of geese and the observed weather data for this period were applied for this validation. The model results show a close match between the observed and simulated numbers of the three ungulate species, although Heck cattle are overestimated by the model, whereas red deer are underestimated (Fig. 2).

**Fig. 2.**
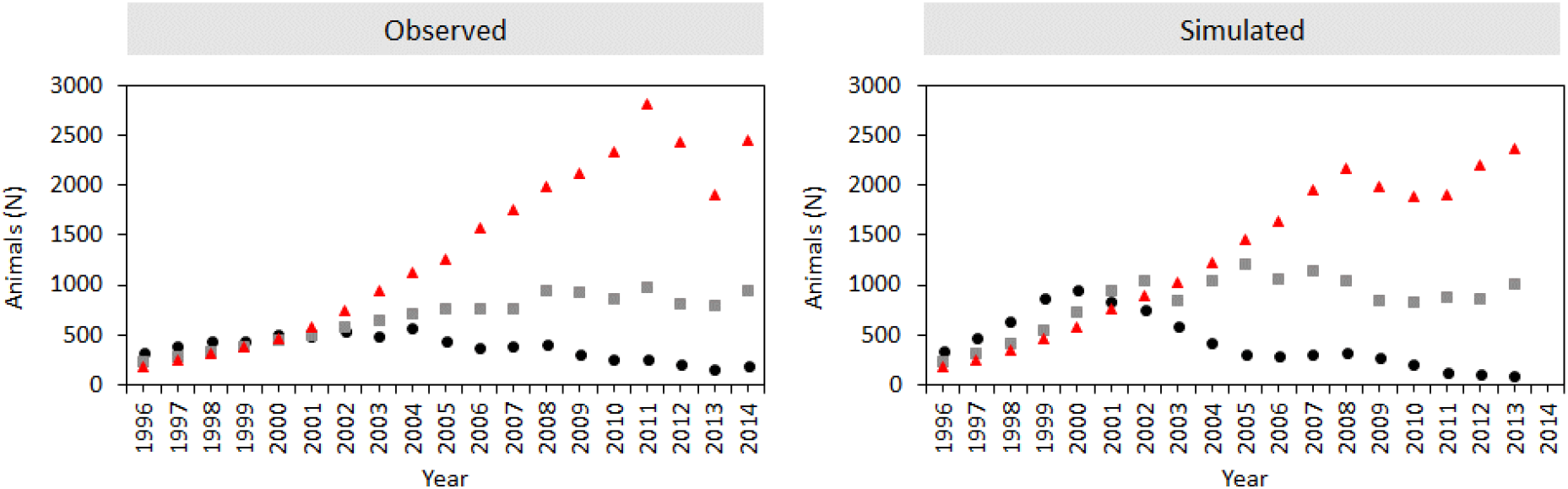
Model versus data comparison with respect to dynamics in numbers of ungulates. Left graph shows the observed numbers of animals of 1 year and older on May 1 of each year; right graph shows the results of the model. Black circles are Heck cattle; grey squares are Konik horses; red triangles are Red deer.

The observed decrease in the number of Heck cattle over time is most likely a result of competition among herbivores, as the sward height of the grasslands in the OVP area decreased below the minimum grazing height for cattle (Cornelissen et al., 2014c). The minimum grazing height for cattle was set at 5 cm (Menard et al., 2002). Konik horses, red deer, and geese can graze more efficiently on swards below 5 cm than cattle (Menard et al., 2002). The minimum grazing height for horses and red deer was set at 2 cm. For geese, a minimum grazing height of 1 cm was used (Durand et al., 2003; Cope et al., 2005). It is assumed that geese, as specialist grazers, only graze on grasses and not on any other plant species in the herb layer (e.g., Aerts et al., 1996; Owen, 1979).

### Model runs

Several scenario analyses were conducted to address the research questions posed in this study. These scenarios included:

a. *Variable versus Constant Weather*: This scenario assessed the effects of weather variability. The temperature series for the variable weather scenario was based on weather time series for the period 1901–2013. For the constant weather scenario, monthly averages from 1901-2013 were used for temperature, incoming radiation, snow cover, and the duration of the growing season, based on the variable weather series.
b. *High Geese Density versus No Geese*: This scenario evaluated the impact of geese presence. The high geese densities were comparable to the numbers that visited the OVP area during 2009-2013.
c. *All Ungulate Species versus No Ungulate Species*: This scenario examined the effects of ungulates on the opportunities for woody species to establish and grow.

We used observed animal numbers and the vegetation map from 1996 as the starting point for our model runs. From that year on, the entire area was grazed by the ungulates.

## Results

The ungulate populations were influenced by both weather and geese (Fig. 3). Weather variability resulted in higher maximum and lower minimum population numbers, leading to increased fluctuations compared to the constant weather scenarios (Fig. 3). Declines in animal numbers were closely linked to severe winters. Geese contributed to decreased ungulate numbers and heightened absolute fluctuations.

**Fig. 3.**
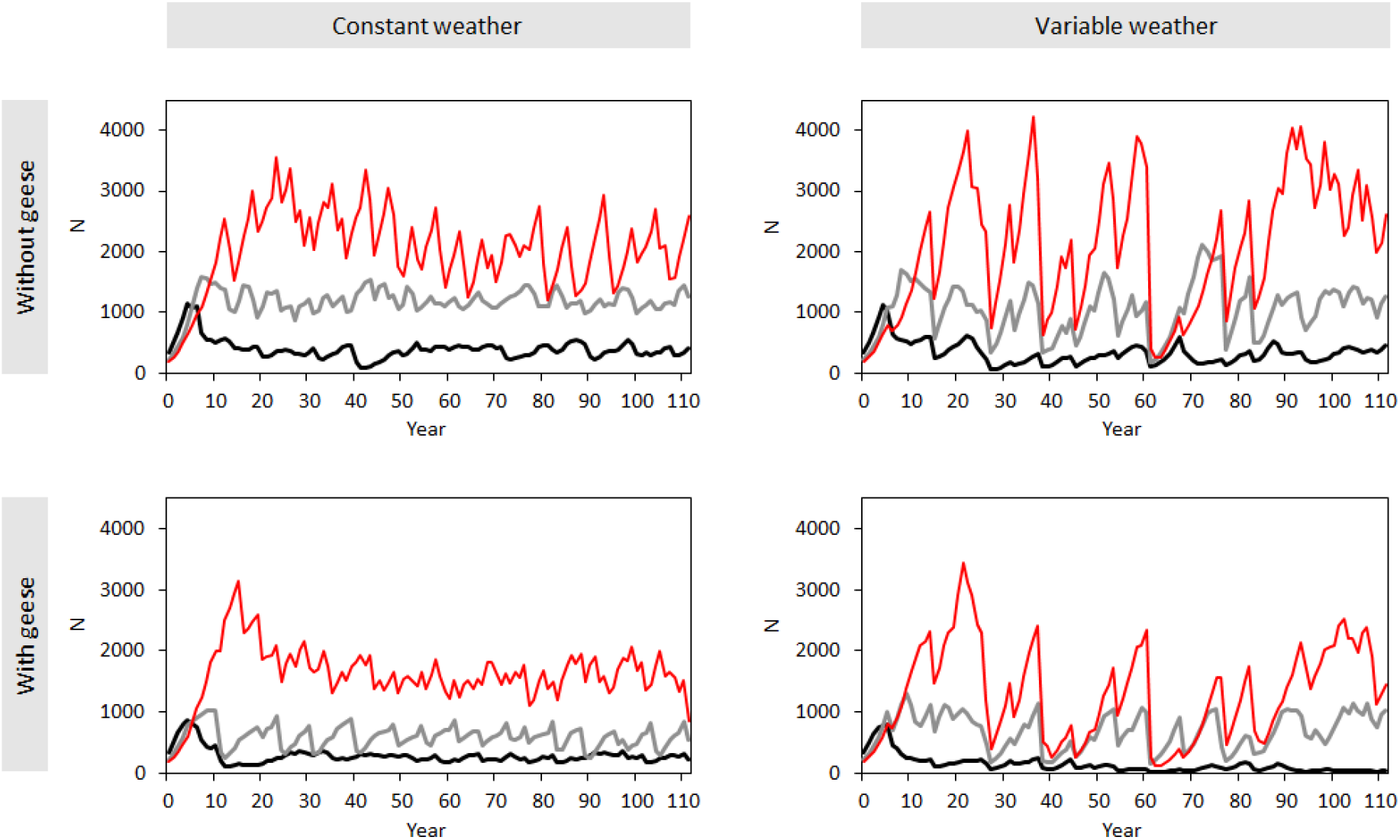
Population dynamics in numbers of the populations of Heck cattle (black line), Konik horse (grey line) and Red deer (red line) in the scenarios with constant and variable weather, and without and with geese.

Weather and geese also impacted relative fluctuations (Fig. 4). The relative increase and decrease— defined as the annual population change expressed as a percentage of the population at the year’s start—were significantly higher in the variable weather scenarios. This effect was also observed in scenarios with geese, though the impact of geese was less pronounced than that of weather variability.

**Fig. 4.**
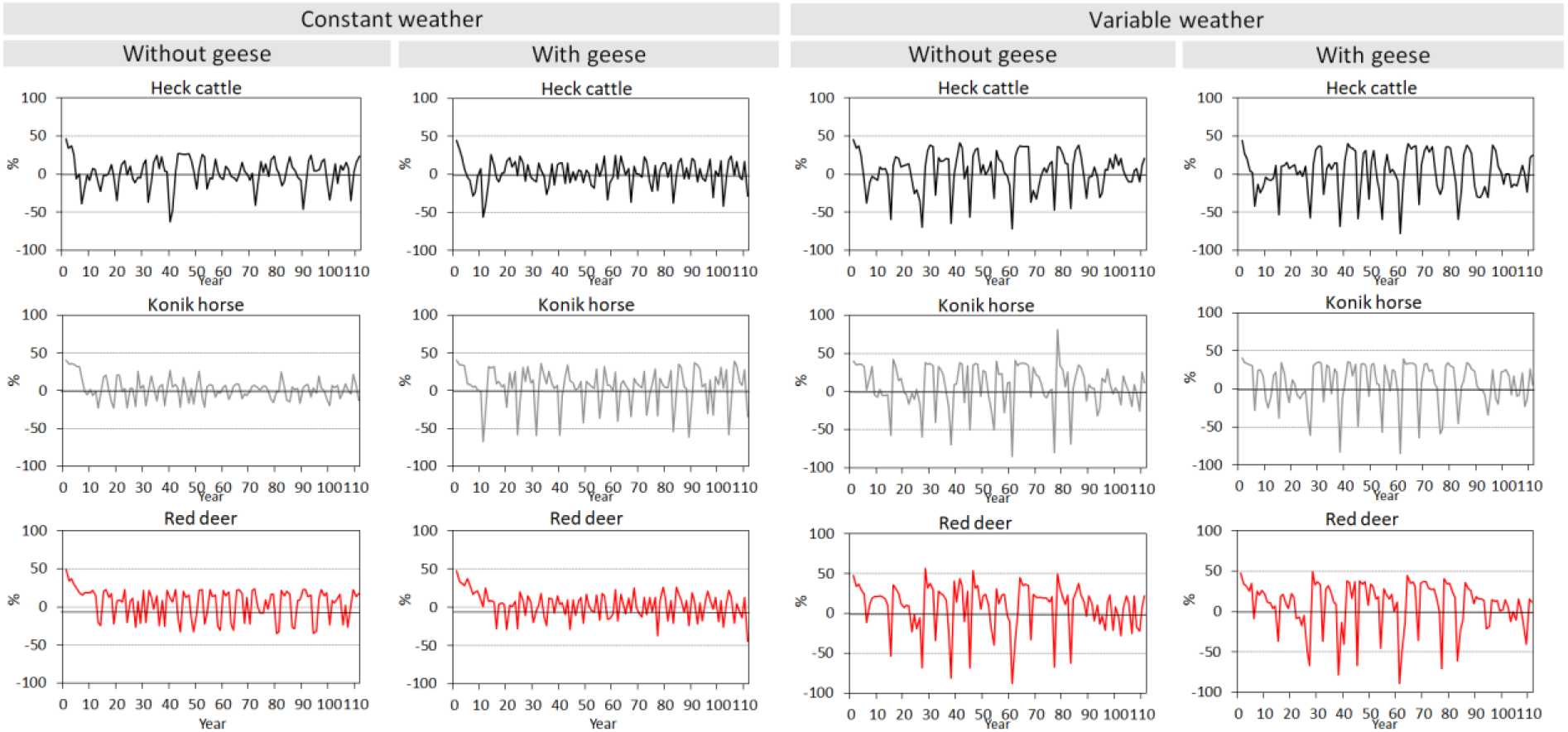
Relative increase and decrease of the populations of Heck cattle, Konik horse and Red deer in the scenarios with constant and variable weather, and without and with geese. The increase or decrease is the change of the population number over one year given as a percentage of the population number at the beginning of that year.

Despite these influences, the three ungulate species coexisted in all weather and geese scenarios over the 110-year period (Fig. 3). Although the number of Heck cattle occasionally dropped very low in the variable weather scenario with high geese numbers, they did not become extinct due to the calibrated effect of winter temperature on herbivore mortality.

Weather and geese influence the development of woody species both directly and indirectly (Figs. 5 and 6). In scenarios without ungulates (Fig. 5), weather variability and the absence of geese resulted in slightly higher cover of woody species compared to scenarios with constant weather and the presence of geese. In these scenarios without herbivores, the area was not completely covered by woody species because the tall growth of herbs and grasses hindered the establishment of the woody species used in the model.

**Fig. 5.**
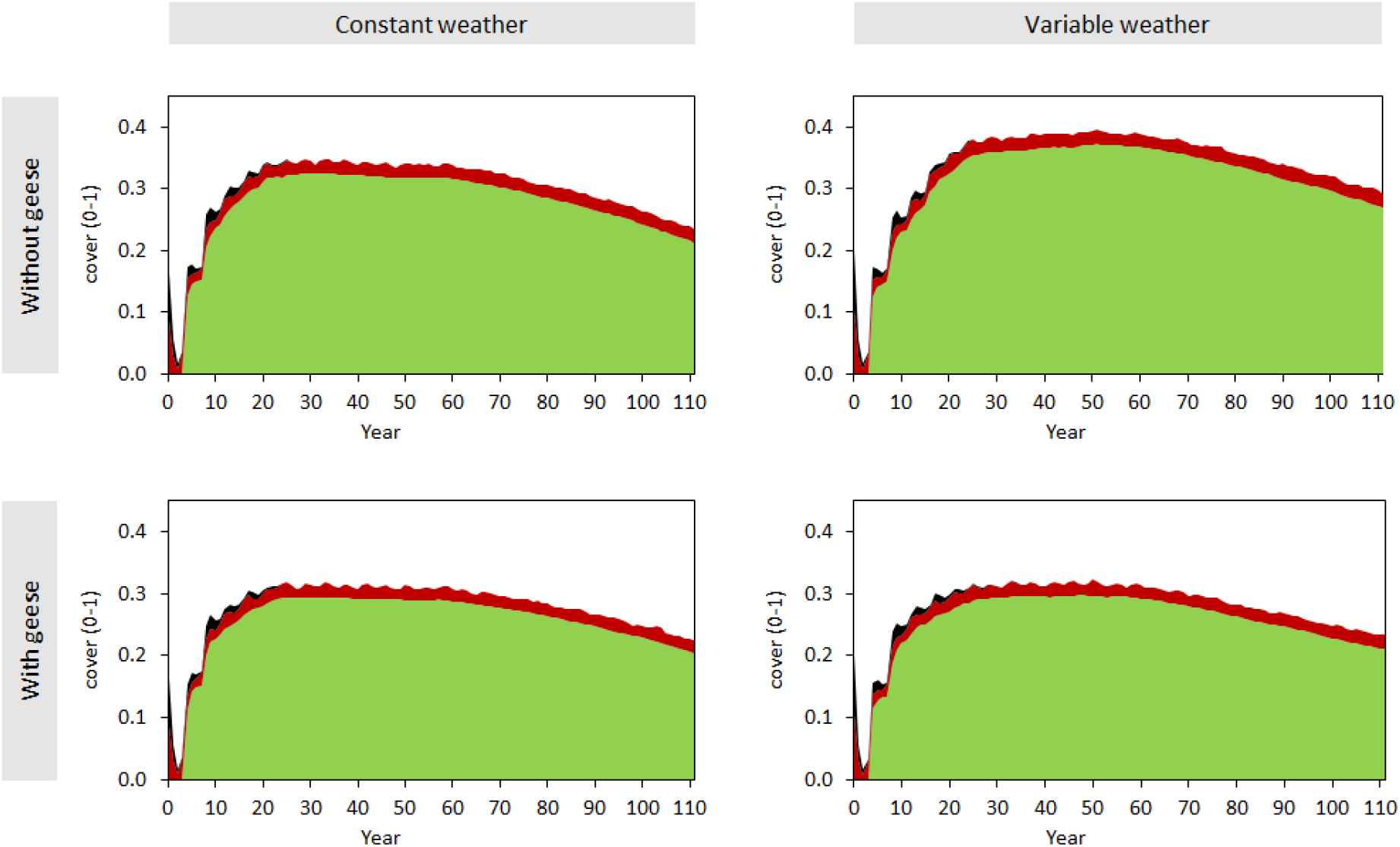
Dynamics of hawthorn (green), willow (red) and elder (black), exceeding 1.5m in height in the scenario without the assemblage of Heck cattle, Konik horses and Red deer, for constant and variable weather and without and with geese. Cover is presented as a proportion between 0-1.

**Fig. 6.**
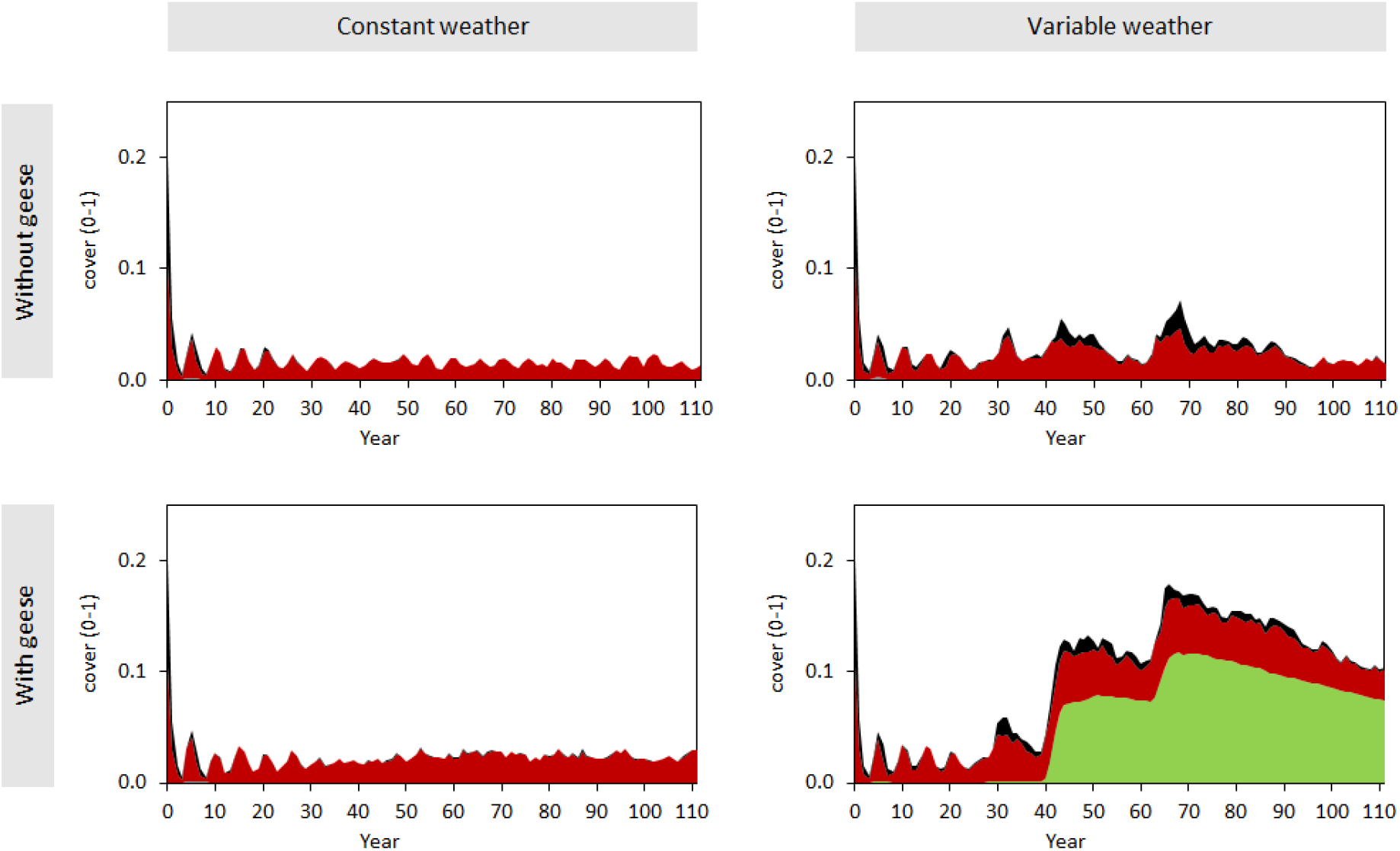
Dynamics of hawthorn (green), willow (red) and elder (black), exceeding 1.5 m in height in the scenario with the assemblage of Heck cattle, Konik horses and Red deer (see fig. 2), for constant and variable weather and without and with geese. Cover is presented as a proportion between 0-1.

Since weather and geese impact ungulate numbers (Fig. 3), both factors also indirectly influence the development of woody species through their effect on ungulate populations. Generally, the indirect effects of ungulates on the development of woody species are much more significant than the direct effects of weather and geese (Figs. 5 and 6).

In scenarios with all ungulates present (Fig. 6), the greatest opportunities for the establishment of woody species occurred with variable weather and the presence of geese, while the least opportunities arose with constant weather and no geese. The thorny shrub hawthorn only established under conditions of variable weather and the presence of geese; elder established only with variable weather, and willow established under all conditions. In the scenario with variable weather and geese, hawthorn established twice—after two periods with the most severe winters—whereas the other two species established more frequently. Once established, hawthorn began to dominate the woody vegetation. The establishment of hawthorn corresponded with a total ungulate number of fewer than 800 animals, which persisted for more than four consecutive years (Fig. 3).

## Discussion

Modeling necessitates balancing generality, realism, and accuracy (Levins, 1966). Levins argues that a model cannot excel in all three aspects simultaneously; at best, two can be optimized, albeit at the cost of the third. In our study, we prioritize realism and accuracy over generality. Therefore, the model must undergo testing and potentially undergo adjustments when applied to new areas, such as adding or removing certain processes. Quantitatively, our findings do not directly apply to other nature reserves. However, qualitatively, we believe that insights gained from the FORSPACE model in the Oostvaardersplassen nature reserve are relevant to other reserves facing similar management challenges. Below, we justify this assertion according to the structure of our research questions for this study:

A. The model results demonstrate that weather variability disrupts ungulate population numbers and influences long-term population dynamics within an ungulate assemblage. Significant reductions in animal numbers lead to increased food availability per capita, fostering opportunities for coexistence. Our expectation was that reduced environmental variability would disadvantage cattle, as they prefer sward heights between 9-16 cm (Menard et al., 2002), whereas Konik horses, red deer, and geese efficiently graze on swards below 5 cm (Menard et al., 2002). High populations of red deer and Konik horses lower grassland sward heights to levels unsuitable for cattle grazing. Even under stable weather conditions, our findings indicate that coexistence among all three ungulate species is feasible. Thus, we conclude that regular weather variability is likely sufficient to prevent competitive exclusion among ungulates, enabling their coexistence.
B. Geese significantly reduced ungulate populations (see Fig. 3). By intensively foraging on regrowth in the short grazed grasslands during winter and spring, the large population of geese maintains a very short sward (<2 cm). Consequently, geese pose strong competition for ungulates during periods of low net primary production. Our findings indicate that the coexistence of the three ungulate species is feasible both in the presence and absence of geese, and under varying weather conditions. However, the presence of cattle is less likely when geese numbers are high and weather variability increases.
C. According to the model results, thorny shrubs can establish and persist in the vegetation over several decades only under conditions of variable weather and high geese populations (see Fig. 6). These factors lead to significant declines in ungulate numbers over multiple years, facilitating the encroachment of thorny shrubs. It is noteworthy that we calibrated the effect of freezing temperatures on maintenance respiration rates to ensure that even the harshest winters do not cause local extinctions of any ungulate species. Therefore, our findings suggest that we may have underestimated, rather than overestimated, the impact of environmental variability on ungulate population dynamics. Despite potential underestimations in winter mortality, our study underscores that combinations of weather variability and high geese densities create favourable conditions for thorny shrub establishment. If freezing temperatures exert stronger effects than simulated, additional opportunities for thorny shrub expansion would likely arise.

The simulated findings presented above are supported by empirical evidence. Cornelissen et al. (2014a, b) demonstrated that in the Oostvaardersplassen (OVP) area, ungulates can transform woody vegetation into grasslands, while woody species only establish at low herbivore densities (<0.5 animals ha-1). Smit et al. (2015) concluded that ungulates can foster wood-pasture landscapes as long as grazing refuges are available. However, high ungulate numbers lead to the absence of these refuges (Cornelissen et al., 2014b). Many other studies also report negative impacts of high herbivore densities on woody species establishment (see Gill, 2006). Our findings align with these studies, indicating that effective wood-pasture cycling is hindered when ungulate numbers remain consistently high without periods of very low densities due to weather fluctuations, diseases, or other factors.

Apart from weather effects, our study highlights the necessity of high geese populations for maintaining low ungulate densities. Small herbivores like rabbits also influence woody species establishment through direct effects such as browsing or debarking (e.g., Kuiters and Slim, 2003; Bakker et al., 2004). In contrast, geese indirectly affect woody species through competition by keeping swards very short. Geese can damage young seedlings in grazed grasslands by trampling or consuming young leaves but do not browse or debark older seedlings or saplings. They prefer highly nutritious young grass leaves over less nutritious woody species, particularly in short grasses, reducing their preference for woody plants. Furthermore, in highly productive areas, geese cannot maintain short swards without ungulates, causing vegetation height to increase within a year (Smit et al., 2015). Vulink (2001) demonstrated that geese prefer intensively grazed, highly nutritious, short swards. As taller swards are less nutritious, they become less attractive to geese, which avoid them.

The general patterns identified are as follows:

i. Ungulates in the Oostvaardersplassen (OVP) facilitate food availability for smaller herbivores, such as geese, by maintaining extensive areas of high-quality shortgrass vegetation. Conversely, smaller herbivores outcompete larger ones, such as cattle in the OVP case, by reducing the availability of shared food sources to a point where survival becomes unsustainable for the larger ungulates. This pattern is expected to hold true regardless of the size or productivity of the nature reserve. However, in smaller or less productive reserves, significant declines in ungulate populations can have a disproportionate impact on their survival compared to larger or more productive reserves. Unregulated populations in smaller reserves tend to be smaller and more vulnerable to declines that may prevent reproduction, particularly affecting sexually dimorphic species like cattle and red deer where male mortality is higher under conditions of food scarcity or severe winters (Clutton-Brock et al., 1982; Georgiadis, 1985; Toïgo and Gaillard, 2003).
ii. Spatial heterogeneity in vegetation can arise from occasional collapses in ungulate densities, allowing shrubs and trees to establish and dominate for extended periods, thereby supporting the validity of the wood-pasture hypothesis even in highly nutrient-rich and productive reserves. However, such opportunities are rare, as observed twice in a 110-year simulation in our case study. In less productive areas, we anticipate more frequent opportunities for establishment under similar weather conditions. This expectation stems from lower food quality per capita, leading to reduced body condition and higher ungulate mortality (Cornelissen and Vulink, 2015).
iii. iii. In nutrient-rich and highly productive areas with ungulates, vegetation tends towards a predominantly open landscape with short grazed swards, occasionally allowing for the establishment of spiny shrub species. Such reserves, managed without ungulate population control and lacking large predators, can support large populations of a few species that favour extensive, short-grass habitats, including migratory birds like geese, ducks, lapwings, and plovers. These areas prioritize the proliferation of a few dominant species (“biomassality”) over maximizing species diversity on a smaller scale. While both biomassality and species diversity contribute to biodiversity, the former is not universally considered as crucial as species diversity in biodiversity conservation efforts.

## Notes

### Competing Interest Statement

The authors have declared no competing interest.

## References

Bugmann, H. 2001. A review of forest gap models Climatic Change. 51:1573–1480.

Bugmann, H.P.J.W. 2003. Forest-Ungulate Interactions: Monitoring, Modeling and Management. Journal of Nature Conservation

Caro, T. and P. Sherman. 2009. Rewilding can cause rather than solve ecological problems. Nature. 462:985.

Cornelissen, P., J. Bokdam, K. Sykora and F. Berendse. 2014. Effects of ungulates on wood pasture dynamics in a European wetland system. Basic and Applied Ecology. 15:396–406.

Cornelissen, P. and J.T. Vulink. 2015. Density-dependent diet selection and body condition of cattle and horses in heterogeneous landscapes. Applied Animal Behaviour and Science. 163:28–38.

Coulson, T., E.A. Catchpole, S.D. Albon, B.J.T. Morgan, J.M. Pemberton, C.-B. T.H., M.J. Crawley and B.T. Grenfell. 2001. Age, sex, density, winter weather, and populations crashes in Soay sheep. Science. 292:1528–1531.

Fontes, L.B. J.D.;Bugmann, H.;Van Oijen, M.;Gracia, C.;Kramer, K.;Lindner, M.;Rötzer, T.;Skovsgaard, J.P. 2010. Models for supporting forest management in a changing environment. Forest Systems. 3:8–29.

Groot Bruinderink, G.W.T.A., D.R. Lammertsma, K. Kramer, J.M. Baveco, A.T. Kuiters, P. Cornelissen, J.T. Vulink, H.H.T. Prins, S.E.v. Wieren, F.E.d. Roder and V. Wigbels. 1998. Draagkracht van de Oostvaardersplassen voor grote herbivoren. Deel 1 Concept en beschrijving van het model. Intern rapport IBN-DLO, Wageningen.

Hodder, K.H. and J.M. Bullock. 2009. Really wild? Naturalistic grazing in modern landscapes. British Wildlife. 20:37–43.

Hopcraft, J.G.C., H. Olff and A.R.E. Sinclair. 2010. Herbivores, resources and risks: alternating regulation along primary environmental gradients in savannas. Trends in Ecology & Evolution. 25:119–128.

Huynh, H.M. 2011. Pleistocene re-wilding is unsound conservation practice. Bioessays. 33:100–102.

Jackson, S.T. and R.J. Hobbs. 2009. Ecological restoration in the light of ecological history. Science. 325:567–569.

Kirby, K.J., R.C. Thomas, R.S. Key, I.F.G. McLean and N. Hodgetts. 1995. Pasture-woodland and its conservation in Britain. Biological Journal of the Linnean Society. 56:135–153.

Kramer, K., G.W.T.A.G. Bruinderink and H.H.T. Prins. 2006. Spatial interactions between ungulate herbivory and forest management. Forest Ecology and Management. 226:238–247.

Kramer, K., T.A. Groen and S.E. Van Wieren. 2003. The interacting effects of ungulates and fire on forest dynamics: an analysis using the model FORSPACE. Forest Ecology and Management. 181:205–222.

Kramer, K., H. Baveco, R.J. Bijlsma, A.P.P.M. Clerkx, J. Dam, J. van Goethem, T.A. Groen, G.W.T.A. Groot Bruinderink, I.T.M. Jorritsma, J. Kalkhoven, A.T. Kuiters, D. Lammertsma, R.A. Prins, M. Sanders, R. Wegman, S.E. van Wieren, S. Wijdeven and R. van der Wijngaart. 2001. Landscape forming processes and diversity of forested landscapes: description and application of the model FORSPACE. Alterra, Wageningen, p 168

Levins, R. 1966. The strategy of model building in population biology. Americal Scientist. 54:421–431.

Menard, C., P. Duncan, G. Fleurance, J.-Y. Georges and M. Lila. 2002. Comparative foraging and nutrition of horses and cattle in European wetlands. Journal of Applied Ecology. 39:120–133.

Navarro, L.M. and H.M. Pereira. 2012. Rewilding abandoned landscapes in Europe. Ecosystems. 15:900–912.

Olff, H., F.W.M. Vera, J. Bokdam, E.S. Bakker, J.M. Gleichman, K. de Maeyer and R. Smit. 1999a. Shifting Mosaics in Grazed Woodlands Driven by the Alternation of Plant Facilitation and Competition. Plant Biology. 1:127–137.

Olff, H., F.W.M. Vera, J. Bokdam, E.S. Bakker, J.M. Gleichman, K.d. Maeyer and R. Smit. 1999b. Shifting mosaics in grazed woodlands driven by the alternation of plant facilitation and competition. Plant Biology. 1:127–137.

Putman, R.J. 1996. Competition & resource partitioning in temperate ungulate assemblies. Chapman & Hall, London.

Rey Benayas, J.M., A.C. Newton, A. Diaz and J.M. Bullock. 2009. Enhancement of biodiversity and ecosystem services by ecological restoration: a meta-analysis. Science. 325: 1121–1124.

Schippers, P., A.J.A. van Teeffelen, J. Verboom, C.C. Vos, K. Kramer and M.F. WallisDeVries. 2014. The impact of ungulates on woodland–grassland dynamics in fragmented landscapes: The role of spatial configuration and disturbance. Ecological Complexity. 17:20–31.

Sinclair, A.R.E., S. Mduma and J.S. Brashares. 2003. Patterns of predation in a diverse predator-prey system. Nature. 425:288–290.

Svenning, J.-C. 2002. A review of natural vegetation openness in north-western Europe. Biological Conservation. 104:133–148.

Vera, F.W.M. 2000. Grazing ecology and forest history. CAB International, Oxford, UK.

WallisDeVries, M.F., Bakker, J.P., Van Wieren, S.E. 1998. Grazing and Conservation Management. Spriner. Berlin, Germany.

Wesseling, C.G., Karssenberg, D., Van Deursen, W.P.A. and P.A. Burrough. 1996. Integrating dynamic environmental models in GIS: the development of a Dynamic Modelling language. Transactions in GIS. 1: 40–48

